# PSMC2 and CXCL8-Modulated Four Critical Gene Biomarkers and Druggable and Vaccinable Targets for Colorectal Cancer

**DOI:** 10.1101/2022.11.15.516622

**Authors:** Yongjun Liu, Yuqing Xu, Xiaoxing Li, Mengke Chen, Xueqin Wang, Ning Zhang, Xiaofei Zhang, Wei Zheng, Heping Zhang, Zhengjun Zhang

## Abstract

Transcriptomic studies have reported numerous differentially expressed genes in colorectal carcinoma (CRC) versus noncancerous tissues. Given the large number of genes identified, it is unclear which ones are the key genes that drive cancer development. To address the issue, we conducted a large-scale study of eight cohorts with thousands of tumor and nontumor samples, analyzed transcriptomic data, and identified the most miniature set of differentially expressed genes (DEGs) that can nearly perfectly describe the overall features of CRC at the genomic level. The analytical framework was built on a recently proven powerful max-linear competing risk factor model. We first analyzed six public transcriptomic datasets and identified four critical DEGs (i.e., CXCL8, PSMC2, APP, and SLC20A1) with nearly perfect (close to 100%) predictive power. The findings were further validated in a newly collected Chinese cohort and another public dataset. Among the four DEGs, PSMC2 and CXCL8 appeared to play a central role, and CXCL8 alone could serve as a biomarker for early-stage CRC. They rise as druggable and vaccinable targets for CRC. This work represents a pioneering effort to identify critical colorectal-specific genes and their interactions that have not been discovered in previous endeavors.

**Simple Summary:** Human knowledge of cancer is still limited. There don’t exist reliable genomic biomarkers for cancer diagnosis, and truly functional and druggable genomic (gene) targets haven’t been reported. One of the main reasons is due to lack of powerful discovery tools to discover the best possible and accurate miniature set of genes to fight against the cancer war. Our research was motivated by such an urgent need, and we hope our findings can fill up gaps in the literature and medical practice. We focus on colorectal cancers in this paper.

## 1. Introduction

Colorectal cancer (CRC) is among the top five most commonly diagnosed malignancy globally [1, 2]. Despite surgical resection and recent advances in chemoradiation and immunotherapy, it remains the second leading cause of cancer mortality worldwide [2, 3]. At present, there is a trend of increasing incidence among young adults in the United States and other countries [4–6]. Genetic predisposition plays an important role in carcinogenesis of CRC [2, 6, 7]. The etiology of CRC can be broadly classified into two categories: hereditary or sporadic. Hereditary CRC accounts for 10-15% of the overall incidence and is attributable to mutations in APC or DNA mismatch repair (MMR) genes. Sporadic CRC is more frequent, representing >80% of CRCs, and is characterized by chromosomal instability (CIN), microsatellite instability (MSI), or CpG island methylation [6, 7].

Over the past decades, transcriptomic studies have been performed to unravel molecular mechanisms underlying colorectal carcinogenesis [8–15]. These published studies differ in study designs and analytical approaches. As a result, massive amounts of genomic data were generated, reporting many differentially expressed genes (DEGs) between tumor and nontumor tissues [8–15]. Using a network-based meta-analysis approach, the recent Consensus Molecular Subtypes (CMS) Consortium analyzed CRC expression profiling data, including The Cancer Genome Atlas (TCGA), in conjunction with molecular data on mutations and somatic copy number alterations [16]. The authors described four molecular subtypes, each of which was characterized by distinct expression profiles of oncogenic/tumor suppressive genes and/or pathways, mutation states of particular genes, and MSI [16].

So far, most transcriptomic studies have used traditional analytical approaches, which rely on fold changes of individual genes between tumor and control tissues or pathway enrichment analysis based on current knowledge of genes and biological processes [11, 17–19]. As a result, the number of genes/transcripts reported is huge, and it is uncertain which of them plays central roles in carcinogenesis and cancer identification/classification. In addition, a small number of critical genes that can describe the overall features of CRC at the genomic level might have been missed in early studies due to the analytical methods adopted. Importantly, in traditional models, gene-gene inter-relationships and gene-disease subtype interrelationships were not satisfactorily addressed. Therefore, there is a need to develop novel methodologies to identify critical DEGs with high sensitivity and specificity. Recent advances in the machine learning community have shown great promise for applying new methods to improve cancer identification/classification and have demonstrated superior performance over traditional methods [20–22].

To this end, we sought to identify the parsimonious subset of critical DEGs for CRC using a newly developed machine-learning method. Our study takes advantage of the max-linear competing structure introduced in the most recently developed max-linear competing factor models [21], max-linear regression models [20], and max-linear logistic models [22]. The major difference between the max-linear structured models and the traditional statistical models is that the original linear combination of predictors is replaced by the maximum of several linear combinations of predictors, called competing factors or competing-risk factors (i.e., signatures). The max-linear structure takes into account the interaction and competing relationship among covariates in predicting the outcome variable, which was overlooked in early models. The max-linear competing risk factor models are different from other conventional machine learning methods and deep learning methods (such as random forest, support vector machine, and group LASSO-based method). The max-linear models have been proven to outperform the existing deep learning methods in terms of prediction power under broad data structures [20, 21]. In this work, using the max-linear competing risk factor models, we analyzed seven transcriptomic datasets, including six public datasets and our newly collected dataset in a Chinese population, and discovered four critical DEGs (CXCL8, PSMC2, APP, and SLC20A1) that displayed the highest possible sensitivity/specificity and robustness for CRC. The results are interpretable and reproducible across diverse cohorts/populations. As a result, these genes can serve as reliable genomic biomarkers, and CXCL8 and PSMC2 rise as druggable and vaccinable targets.

## 2. Materials and Methods

### 2.1 Data acquisition and processing

Six publicly available whole-transcriptome datasets were used for discover analyses. The public datasets included two RNA-sequencing datasets from The Cancer Genome Atlas (TCGA), and four mRNA expression chip data of CRC from the Gene Expression Omnibus (GEO) database by using the keywords of “colon cancer” and “Homo sapiens”. Since the primary purpose of our study was to identify critical DEGs for CRC in general, we chose the transcriptome datasets from studies of different platforms (RNA-seq vs. microarray), populations/ethnicities (North American and European Caucasians, Blacks, Chinese, Japanese, and Jewish), and tumor stages. In addition, the relevant clinical and pathological information, such as age, sex, American Joint Committee on Cancer (AJCC) TNM tumor stages and histologic grades of the tumor, was also acquired when available.

The first public dataset was downloaded from the NCI’s Genomic Data Commons (GDC, https://portal.gdc.cancer.gov/). This was an RNA-seq gene expression study performed in the TCGA Colon Cancer (COAD) cohort using the Illumina HiSeq 2000 RNA Sequencing platform [23]. The dataset was published on July 19, 2019 and contained 288 CRC samples and 41 normal controls with 20, 530 genes/transcripts analyzed. Gene expression values were log_2(norm count+1) transformed.

The second public dataset was also obtained from the NCI’s GDC. This was an RNA-seq gene expression study of CRC performed in the TCGA COAD cohort using the Illumina HiSeq 2000 RNA Sequencing platform [23]. The expression values were normalized with log_2(Fragments Per Kilobase of transcript per Million mapped reads (FPKM)+1). The dataset was published on October 13, 2017, containing 471 CRC samples and 41 normal controls with 64,083 genes/transcripts analyzed. There was partial overlap between the first and second datasets: the 41 normal controls were identical. However, gene expression values were scaled in different ways between the two datasets.

The third public dataset (GSE39582) was obtained from a European multicenter study using the Affymetrix Human Genome U133 Plus 2.0 Array platform [11]. This dataset included 566 CRC samples and 19 normal controls with 54, 675 genes/transcripts analyzed [11]. The expression values were derived from log_2(normalized intensity signal).

The fourth public dataset (GSE9348) was obtained from a genome-wide expression profiling study of 82 age-, ethnicity- and tissue-matched Han Chinese CRC patients with healthy controls using the Affymetrix U133 Plus 2 array [24]. All the patients were classified as early-stage CRC (Stage 1/2) [24]. Gene expression values are MAS5-calculated signal intensity.

The fifth dataset (GSE18105) was obtained from a microarray-based genome-wide expression profiling study of 111 samples (77 CRC samples and 17 paired samples from adjacent nontumor tissues) in a Japanese cohort [25]. The patients in this cohort were classified as stages 2/3 CRC [25]. Gene expression values are RMA signal intensity.

The sixth public dataset (GSE41258) was obtained from a microarray-based genome-wide expression profiling study performed in an Israel population including 299 samples (180 primary CRC, 46 polyps, 43 normal colon, 21 liver metastases, and 9 lung metastases) [26]. Data were PLIER normalized batch correct then Lowess normalized signal.

In addition to the above public datasets, we collected a Chinese cohort (the seventh dataset) at Sun Yat-sen University of Medical Sciences, China, for validation analysis. This cohort contained 45 CRC samples and 47 normal controls. Gene expression analysis was performed by real-time quantitative RT-PCR using the TaqMan Gene Expression assays (Applied Biosystems, Inc.), focusing on the four critical DEGs (i.e., CXCL8, PSMC2, SLC20A1, and APP) identified in the public datasets. The patients’ age, sex, TNM tumor stage, and tumor histologic grade were available for analysis. The collection procedures of clinical specimens were approved by the Clinical Research Institution Review Committee and Ethics Review Committee of Sun Yat-sen University of Medical Sciences. The patient provided ‘written’ informed consent before sample collection.

We further tested the identified four genes using the eighth dataset (GSE103512 [52]) and obtained an overall accuracy of 98.55%, a sensitivity of 100%, and a specificity of 91.67%.

Table 1 summarizes the general information of the datasets, including the data sources, experimental platforms, sample sizes, populations, and tumor stages.

**Table 1.**
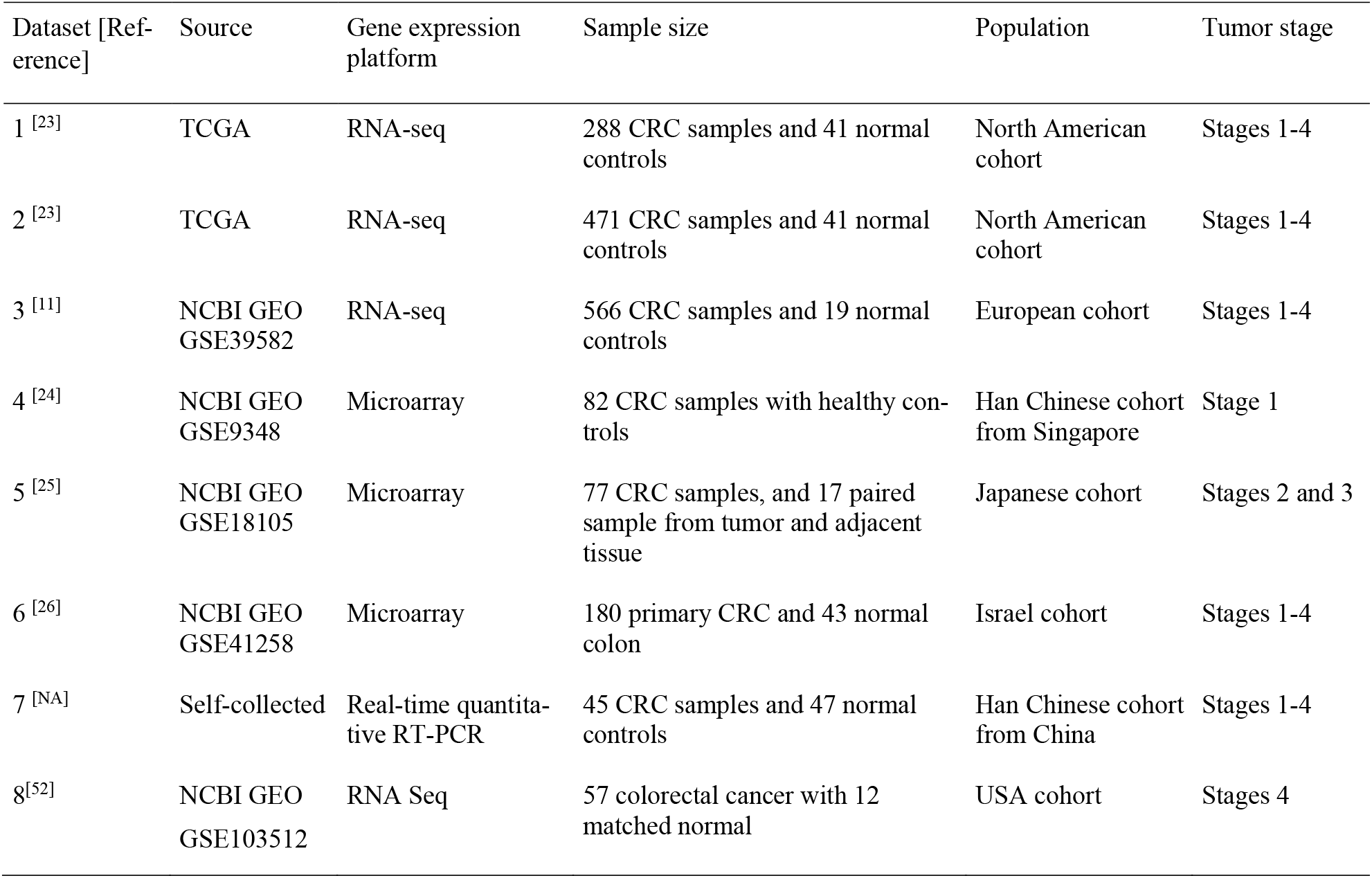
Information of the seven public datasets and one independent Chinese dataset

| Dataset [Ref-<br>erence] | Source | Gene expression<br>platform | Sample size | Population | Tumor stage |
| --- | --- | --- | --- | --- | --- |
| 1 <sup>[23]</sup> | TCGA | RNA-seq | 288 CRC samples and 41 normal controls | North American cohort | Stages 1-4 |
| 2 <sup>[23]</sup> | TCGA | RNA-seq | 471 CRC samples and 41 normal controls | North American cohort | Stages 1-4 |
| 3 <sup>[11]</sup> | NCBI GEO<br>GSE39582 | RNA-seq | 566 CRC samples and 19 normal controls | European cohort | Stages 1-4 |
| 4 <sup>[24]</sup> | NCBI GEO<br>GSE9348 | Microarray | 82 CRC samples with healthy controls | Han Chinese cohort from Singapore | Stage 1 |
| 5 <sup>[25]</sup> | NCBI GEO<br>GSE18105 | Microarray | 77 CRC samples, and 17 paired sample from tumor and adjacent tissue | Japanese cohort | Stages 2 and 3 |
| 6 <sup>[26]</sup> | NCBI GEO<br>GSE41258 | Microarray | 180 primary CRC and 43 normal colon | Israel cohort | Stages 1-4 |
| 7 <sup>[NA]</sup> | Self-collected | Real-time quantitative RT-PCR | 45 CRC samples and 47 normal controls | Han Chinese cohort from China | Stages 1-4 |
| 8<sup>[52]</sup> | NCBI GEO<br>GSE103512 | RNA Seq | 57 colorectal cancer with 12 matched normal | USA cohort | Stages 4 |

### 2.2 Analytical Methodology

We implemented the max-linear logistic regression model to build a competing risk factor classifier for CRC. The competing factor classifier has an advantage over existing models in nonlinear predictions and classifications.

The goal was to select a parsimonious subset of key genes with the highest performance. The theoretical foundation of competing risk factor models was recently described elsewhere [20, 21, 27]. The heterogeneous extension of the max-linear logistic regression was used to consistently study the critical DEGs across the six public datasets and our Chinese cohort. We started with three competing risk factors in the max-linear logistic regression models, with each factor having only three genes randomly drawn from the genes/transcripts in each dataset. Then, a Monte Carlo method with extensive computation was used to find the final model with the best performance of sensitivity and specificity and the smallest number of genes. The basic ideas of competing risk classifiers for heterogeneous populations are described below.

Suppose there are *K* cohorts with their binary (1 for CRC, 0 for CRC free) response variables being ***Y***_(1)_,…, ***Y***_(*K*)_ where

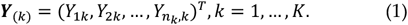

Each of the *Y_ik_*, (*i* = 1,…, *n_k_, k* = 1,…, *K*) may be related to *G* groups of genes

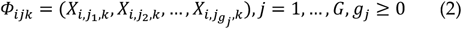

where *i* is the *i*th individual in the sample, *g_j_* is the number ot genes in jth group. The competing (risk) factor classifier for the fcth outcome variable is defined as

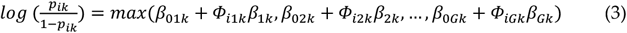

where *β*_0*jk*_’s are intercepts, *Φ_ijk_* is a 1×*g_i_* observed vector, *β_jk_* is a *g_j_* × 1 coefficient vector which characterizes the contribution of each predictor to the outcome variable ***Y***_(*k*)_ in the *j* th group to the risk, and *β*_0*jk*_+*Φ_ijk_β_jk_* is called the *j*th competing risk factor, i.e., *j*th signature.

#### Remark 1

With *β*_0*jk*_ = -∞, *j* = 2,…, *G*, (3) is reduced to the classical logistic regression classifier. It is clear that every component of *β*_0*jk*_ + *Φ_ijk_β_jk_, j* = 1,…, *G* is a risk factor for a patient to have CRC, and the highest risk is from the largest one, i.e., these risk factors compete against each other to win out to make the final effect, i.e., to determine whether or not a patient is CRC. As such, they are called competing (risk) factors. We note that even only one makes the final effect, which does not mean the other risk factors are useless. Because each risk factor represents one potential symptom (subtype) of CRC due to the different combinations of the DEGs, every component risk factor will be useful for clustering a patient into a particular CRC subtype according to the computed risk probability.

The unknown parameters are estimated from

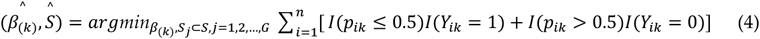

where 0.5 is a probability threshold value that is commonly used in machine learning classifiers, *I*(.) is an indicator function, *p_i_* is defined in Eqn. (3), *S* = {1, 2,…, 54675} is the index set of all genes, *S*_1_ = {1_1_, 1_2_,…, 1_*g*1_}, *S*_2_ = {2_1_,…, 2_*g*2_},…, *S_G_* = {*G*_1_,…, *G_gG_*} are index sets corresponding to (2), and 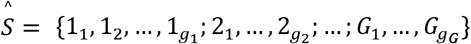 is the final gene set selected in the final classifiers.

To introduce sparsity for both the number of variables (genes) and the number of groups (competing factors, signatures) into the model, the following optimization problem with penalties is considered:

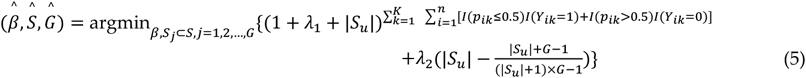

where *S_u_* is the union set of 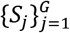, |·| is the cardinality. Tuning parameters *λ*_1_ and *λ*_2_ are both non-negative. 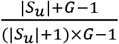 is monotone decreasing in both |*S_u_*| and *G*. Additional properties of this bivariate function was described elsewhere [20].

#### Remark 2

In (2), *X_j_l_,k_1__* and *X_j_l_,k_2__* can be measured under different scales for *k*_1_ ≠ *k_2_* even if they correspond to the same genes (variables), i.e., from heterogeneous populations or cohort studies.

#### Remark 3

(5) is a completely new machine learning classifier with completely different penalization from existing ones, such as LASSO, SCAD, and MCP.

Next, we show a unique theoretical and computational property of the new competing risk factor classifier. The optimization problem (5) is designed to guarantee that, with suitable choices of *λ*_1_ ≥ 0 and *λ*_1_ ≥ 0, the solution of problem (5) will lead to the smallest number of subsets of variables (|*S_u_*|) and the smaller number of signatures (S4) (*G*) while achieving the best possible minimal misclassification rate.

The rationale is as follows:

1. Suppose the underlying best classifier is a ‘perfect classifier’, with 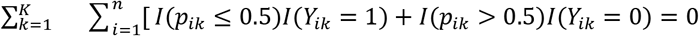 then with this classifier

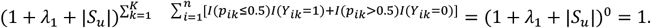 Problem (5) is equivalent to

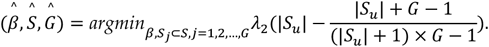 Since |*S_u_*| ≥ 1, 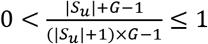, problem (5) is equivalent to first minimiing *S_u_* and then G, which leads to the smallest possible |*S_u_*| and *G*.
2. Suppose the underlying best classifier is not a perfect classifier, with the minimal misclassification number

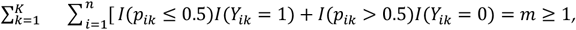

then there exists *λ*_1_ ≥ 0 and *λ*_2_
≥ 0such that

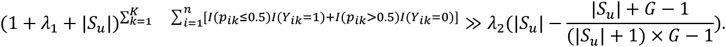 Therefore, problem (5) will first minimize,

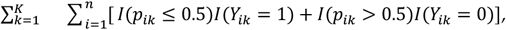

then |*S_u_*|, and finally *G*, which will again lead to the smallest possible |*S_u_*| and *G*.

#### Remark 4.

The S4 property of (5) and its capability to simultaneously classify multiple heterogeneous populations with common variables (genes) make the new competing risk factor classifier different from existing ones.

#### Remark 5.

When *K* = 1 and *λ*_2_ = 0, (5) is equivalent to the classifier introduced by Zhang [22] in a study which identified five critical genes associated with seven Covid-19 subtypes. The details of computational steps were described early [22], and demo Matlab^OR^ codes are publicly available online.

Note that Equation (5) is an integration of integer programming, combinatorial optimization, and continuous optimization. Its computational complexity level is extremely high. In this study, we adopted the following Monte Carlo approach:

1. Randomly selecting a cohort (population), say *k = 1* without loss of generality.
  a. Randomly draw *G* sets of genes with each set having |*S_u_*| genes;
  b. Use any optimization procedures (e.g., Nelder–Mead method, genetic algorithm, simulated annealing) to solve minimizing 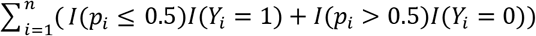.
  c. Repeat the above two steps for *G* = 1, 2, 3 and |*S_u_*| = 1, 2, 3, 4, 5 until an acceptable best solution is reached.
2. Using the genes selected from the *k* = 1 cohort for *k* = 2,…, *K*, repeat the above two steps (b) and (c) for *G* = 1, 2, 3 until an acceptable best solution is reached.

#### Remark 6.

For the data used in this study, a nearly perfect classifier was achieved using the above Monte Carlo approach.

We adopted the following criteria to define critical DEGs:

1. The number of genes should be as small as possible (smaller than 15).
2. This set of genes leads to an overall accuracy of > 95% in at least three different study cohorts with a total number of patients/subjects of at least 1,000.
3. This set of genes leads to an overall 100% accuracy for at least one study cohort with at least 10 subjects.
4. At least one gene functions and shows the same sign (+ or -) in each study cohort.
5. This set of genes should lead to at least 80% accuracy for any cohort with either sensitivity or specificity of >75%.
6. In each competing classifier, the number of genes should be as small as possible, and it must be less than six.
7. The number of competing classifiers should be as small as possible, and without redundancy, i.e., every classifier cannot be replaced.

## 3. Results

### Identification of critical DEGs

We identified four critical DEGs, namely, CXCL8 (C-X-C Motif Chemokine Ligand 8), PSMC2 (Proteasome 26S Subunit, ATPase 2), APP (Amyloid Beta Precursor Protein), and SLC20A1 (Solute Carrier Family 20 Member 1).

### Identification of classifiers based on the four critical DEGs

Each of the competing factors (*CF**_i_, i*** = 1, 2,) is a linear combination of gene expression for a single-digit number of critical genes. The final classifiers were the combined classifiers of the three competing factors, as shown in Table 2. The risk probabilities were calculated using the logistic function of ***exp*** (*Data_**i**_CF_max_*)/(**1 + *exp*** (*Data_**i**_CF_max_*)) for the combined classifiers in each dataset, and of ***exp*** (*Data_**i**_CF_**j**_*)/(**1 + *exp*** (*Data_**i**_CF_**j**_*)) for each individual classifier ***i*** = 1, 2, 3, ***j*** = 1, 2, 3. Data_*i_CF_j_* each can represent one type of gene-CRC relationship and reveal how genes interact with each other. Multiple types (e.g., j=1, 2, 3) of gene expression combinations for a certain patient could predispose to CRC. In other words, they are the competing risk factors for each patient. Data_*i_*CFmax, *i*=1, 2, 3, is the combined maximum of linear competing factors of the *i*th dataset, and the ‘consequence’ of the internal competing over all competing factors.

**Table 2.**
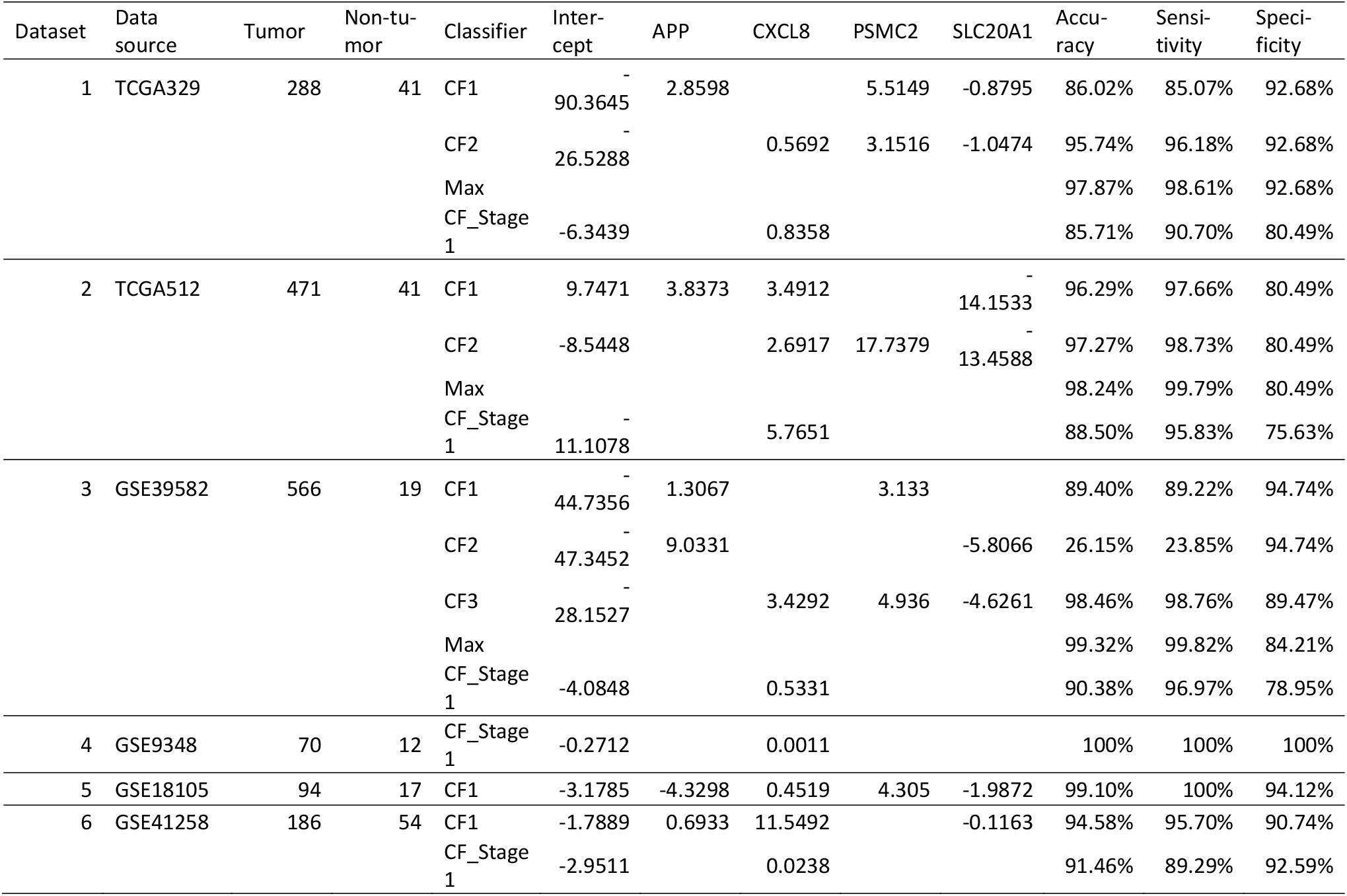
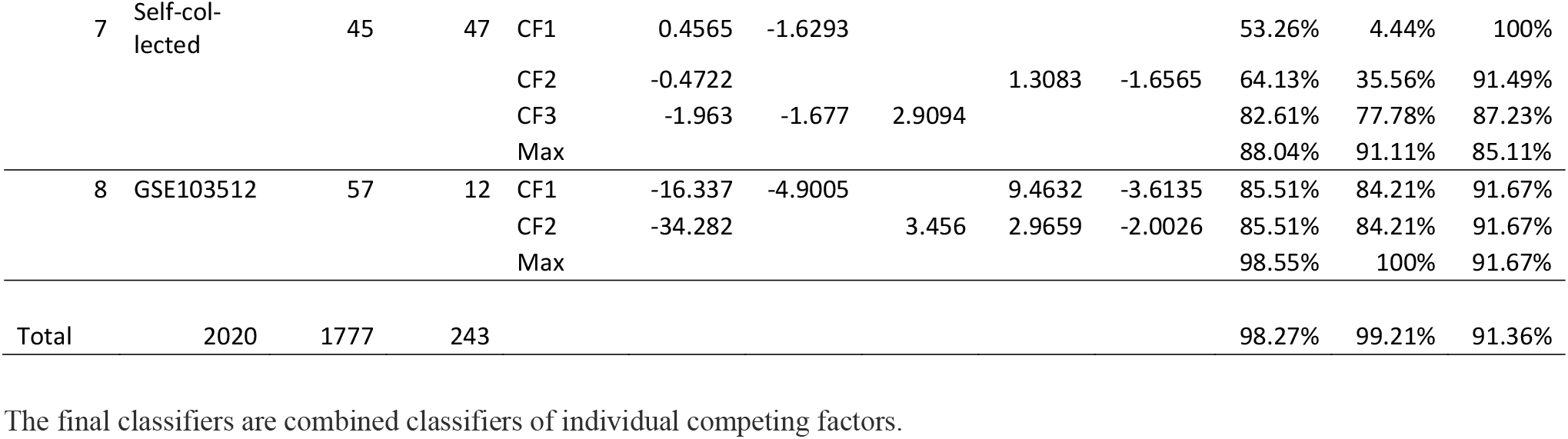
The four critical DEGs and the classifiers identified in the eight datasets

In the first and second datasets, CF1 and CF2 classifiers had a sufficiently high power to identify CRC patients; thus, additional CFs were not needed for cancer identification. In the third and seventh datasets, three classifiers (i.e., CF1, CF2 and CF3) were required to accurately predict the existence of CRC tumors, due to the low sensitivity of CF2 in these two datasets. In the remaining datasets, only one classifier, CF1, is needed for the identification of CRC with high sensitivity and specificity.

For illustration purposes, Figure 1 shows the model-estimated risk probabilities evaluated from the final classifiers in the first three datasets.

**Figure 1.**
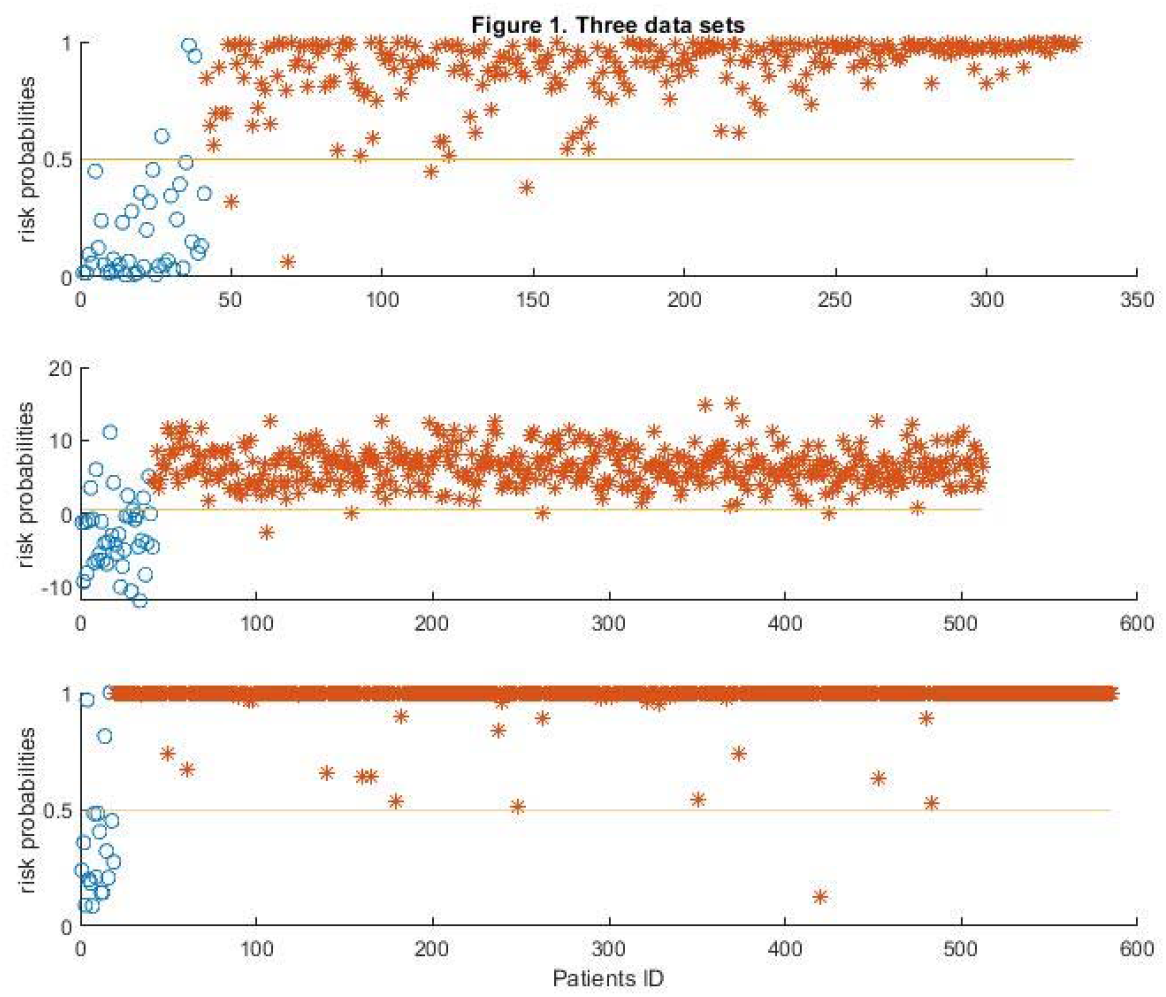
Model-estimated risk probabilities evaluated from the three final classifiers in the first three datasets. Note: Colorectal cancer (CRC) samples are designated by asters, and normal controls are designated by circles. The plot designates the CRC samples by asters and normal controls (NC) by circles. In addition, a 0.5 (50%) horizontal line (probability threshold value) is plotted in each panel.

Figure 2 is a four-dimension plot illustrating the signature patterns defined by each classifier in the third dataset. One can clearly appreciate how the genes form signature patterns in a geometry space. For example, the classifiers identified are used to separate the yellow colors from the blue and green colors. We note that only these four particular genes can show such patterns, i.e., one cannot arbitrarily select three genes to obtain such signature patterns.

**Figure 2.**
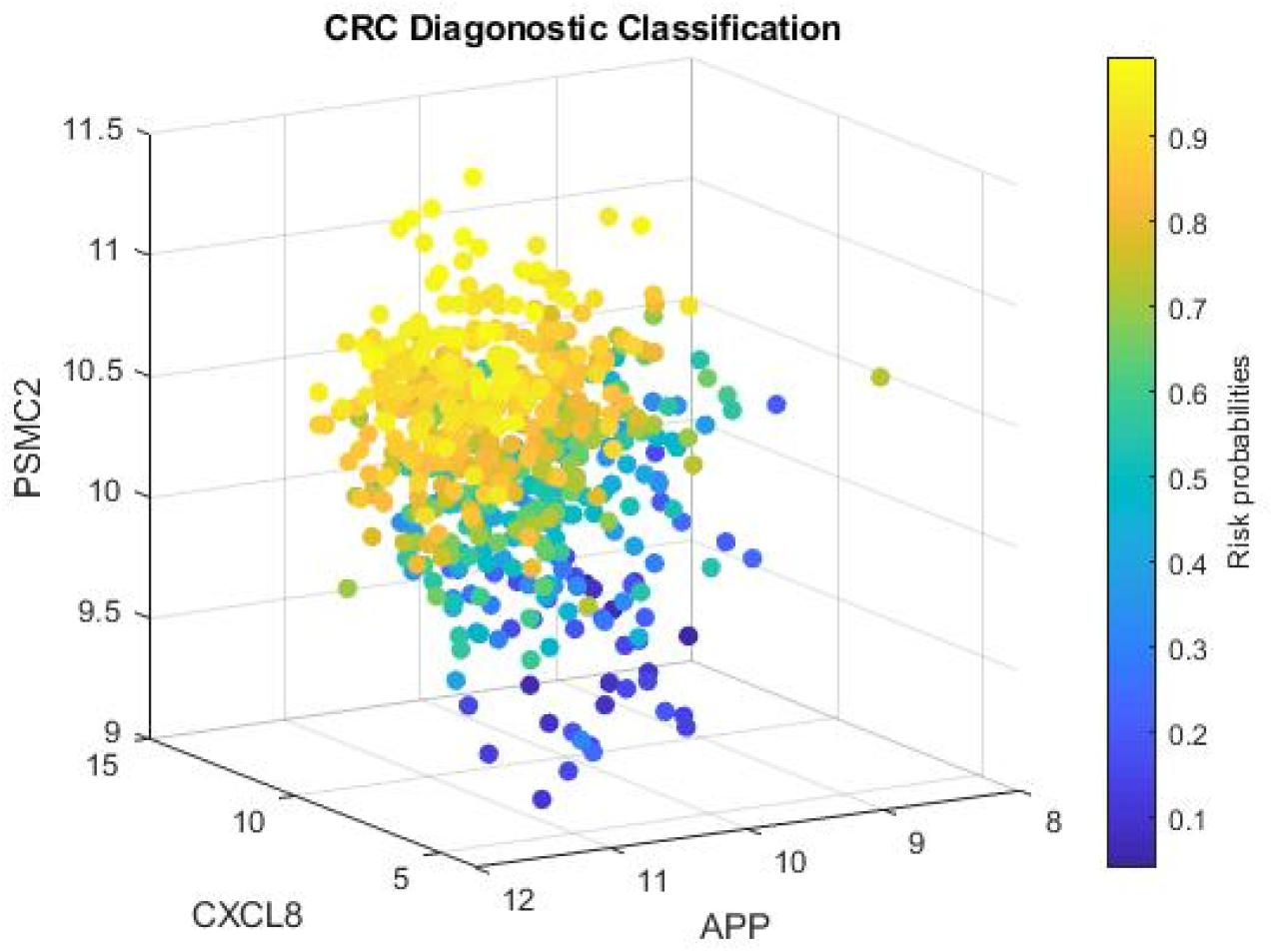
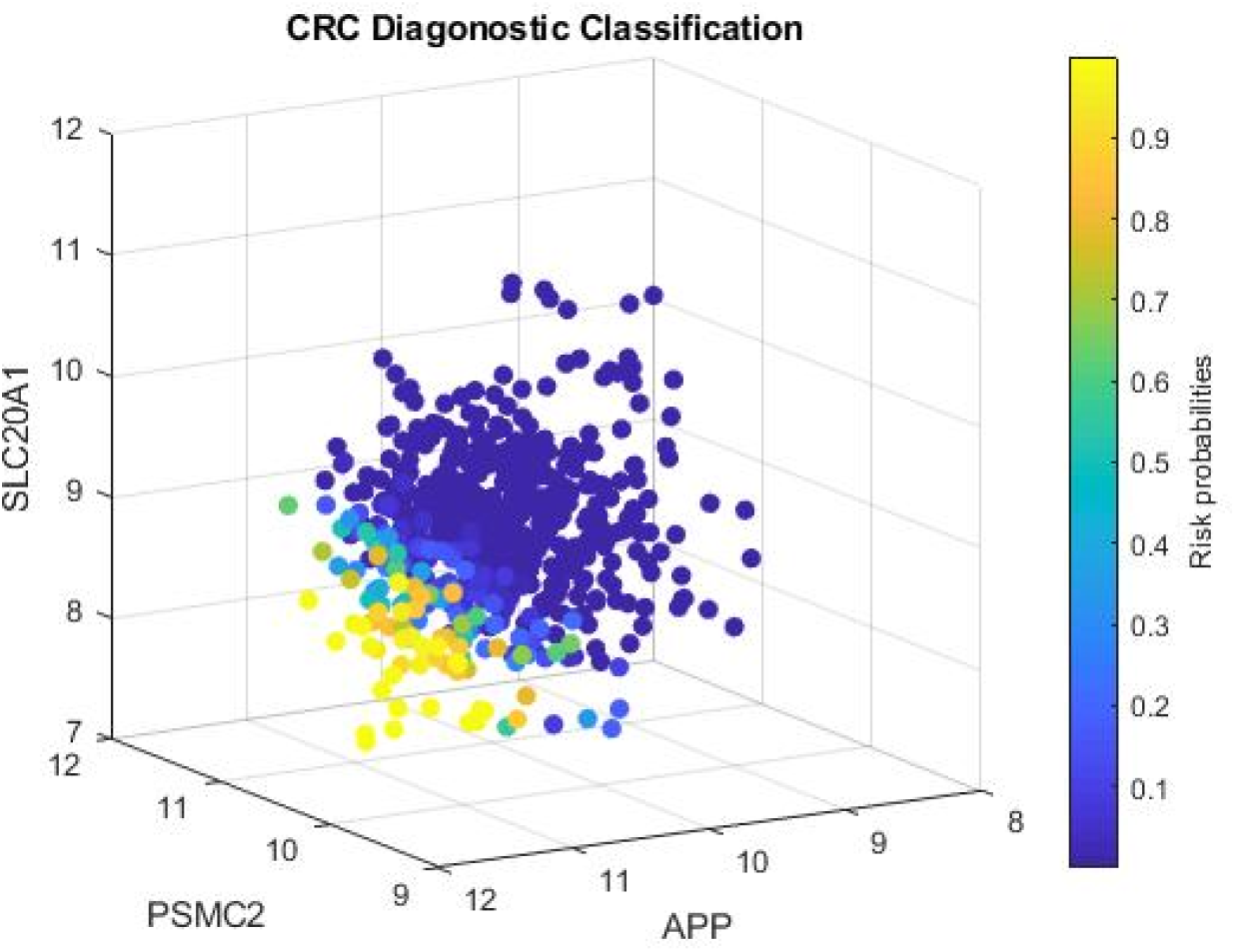
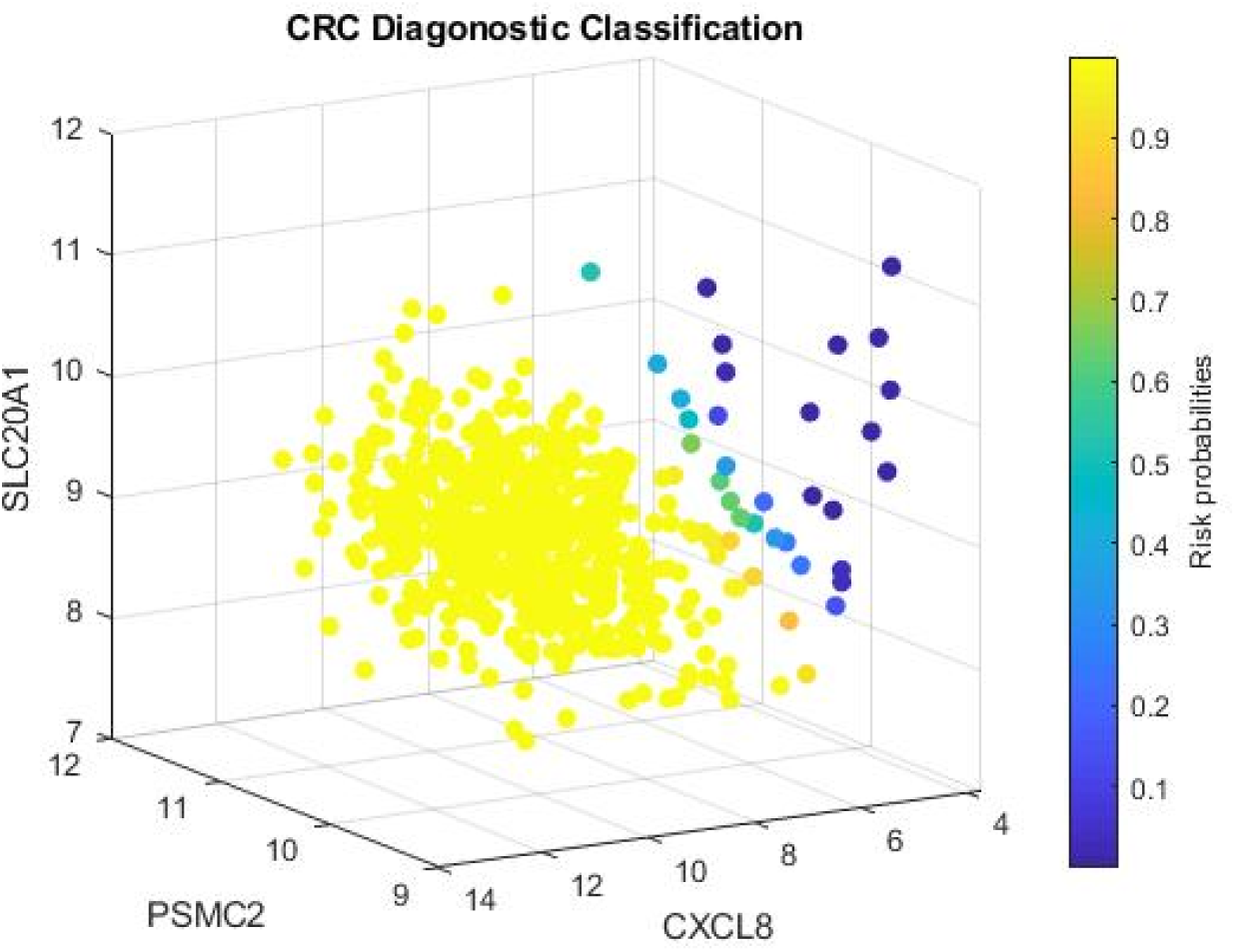
Four-dimensional plots for visualizing risk signature patterns from 3 competing component classifiers in the European cohort (dataset 3) Note: Gene expression values and their combination effects with different strengths are shown in each plot. The fourth dimension represents the risk probabilities. The plot provides a straightforward view of identifying patients with a high risk of CRC.

Figure 3 is a Venn diagram demonstrating patient subgroups classified by the classifiers in the first three cohorts and the validating Chinese cohort (seventh cohort). CRC patients could be classified into three subgroups in the first and second cohorts based on the above classifiers. The first subgroup contained the patients who were only detected by the CF1, the second subgroup contained the patients who were only detected by the CF2, and the third subgroup contained the patients who were detected by CF1 and CF2, simultaneously (Figure 3). In the third cohort, CRC patients were classified into seven subgroups based on the classifiers CF1, CF2, and CF3. In the validating Chinese cohort, patients were classified into seven groups, but some groups were devoid of patients. In the remaining datasets (datasets 4, 5, and 6), patients were not further subclassified as one classifier had sufficient power to identify CRC patients. This figure shows the complexity of disease status as patients fall into different regions from different subgroups with different gene-gene interactions at genomic levels, which can determine the effectiveness of CRC diagnosis, prognosis, and management.

**Figure 3.**
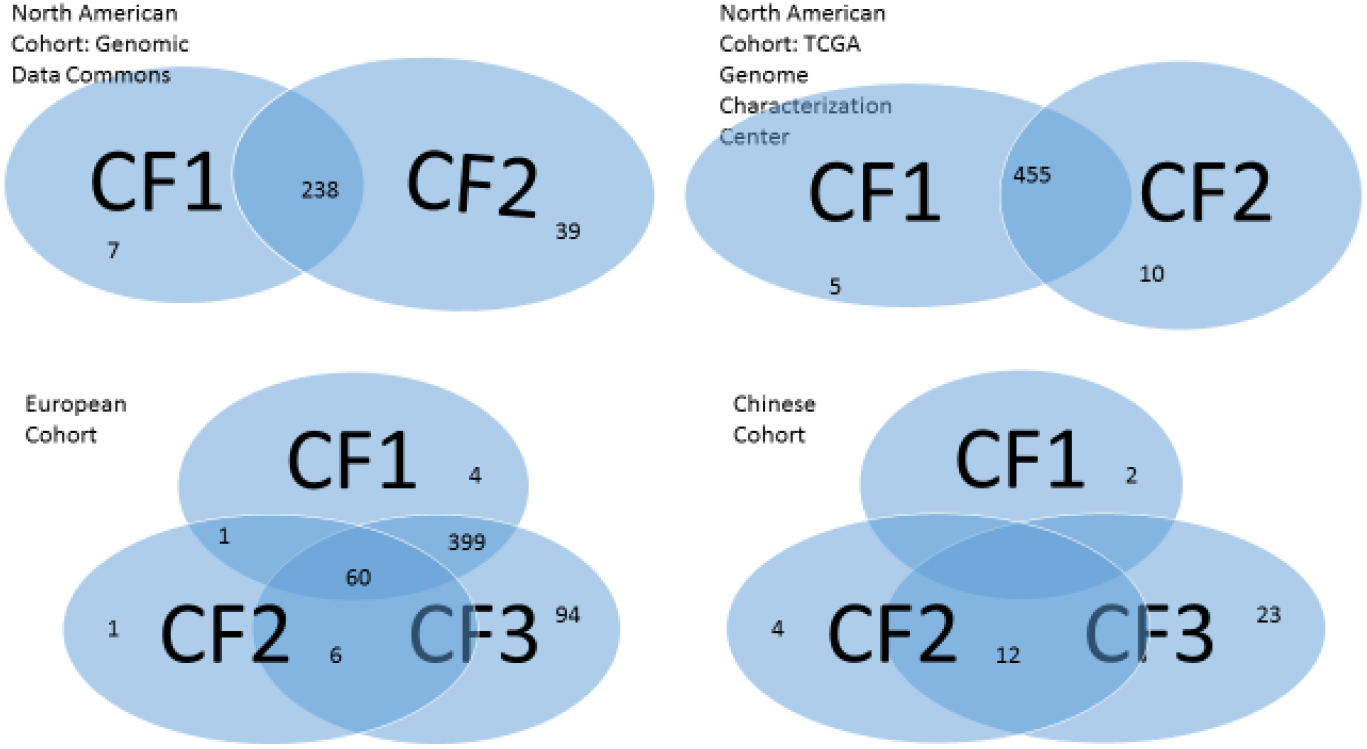
Venn diagrams of subgroups of colorectal cancer patients

As shown in Table 2, the classifier CFmax had a decent performance in differentiating tumor vs. nontumor samples, with an overall sensitivity of >98%, specificity of >80%, and accuracy of >94%. Applying CF1, CF2, and CF3 simultaneously increased the power of cancer detection. Interestingly, in the fourth dataset which included early-stage CRC only in a Han Chinese population, we found the classifier defined by CXCL8 alone could identify CRC with 100% sensitivity/specificity/accuracy. We, therefore, performed focused analyses on early-stage CRC (stage 1) in other datasets. It can be seen CXCL8 alone had high sensitivity (>89%), specificity (>75%), and accuracy (>85%) in the identification of stage 1 CRC in datasets 1, 2, 3, and 6. Since dataset 5 only contained stages, 2 and 3 CRC and dataset 7 contained only 3 cases of stage 1 CRC, further evaluation of stage 1 CRC was not performed in these two datasets.

The risk probability of CRC was determined by the direction and absolute value of the coefficient of the classifiers. A positive coefficient indicated a higher gene expression value was associated with a higher risk probability of CRC. On the contrary, a negative coefficient suggested that a lower gene expression value was associated with a higher risk probability of CRC. For instance, in datasets 1, 2, 3, and 7, decreased APP gene expression was associated with a reduced risk probability of CRC. On the contrary, APP showed the reversed direction in datasets 5 and 7, suggesting different effects of this gene in CRC between Caucasian and Asian populations. In all the datasets, decreased SLC20A1 expression was associated with increased risk probability of CRC, suggesting expression of this gene was suppressed in CRC patients of different populations. Conversely, in all the datasets, decreased CXCL8 and PSMC2 expression was associated with a reduced risk probability of CRC. Different signs of APP in different classifiers suggested that the relationship of a single human gene can be nonlinearly correlated with the disease status, and its interaction with other genes can be positively or negatively correlated.

As shown in Table 2, coefficients associated with CXCL8 and PSMC2 were positive in all the datasets. Considering the heterogeneity of the seven cohorts, it is safe to infer that CXCL8 and PSMC2 were the centers of genegene and gene-subtype interactions. The effects of these classifiers (competing risk factors) were largely modulated by CXCL8 and PSMC2.

For the purpose of illustration, Table 3 shows gene expression values of the four critical DEGs in a small portion of the datasets. The full datasets with original gene expression and computed values are available online.

**Table 3.**
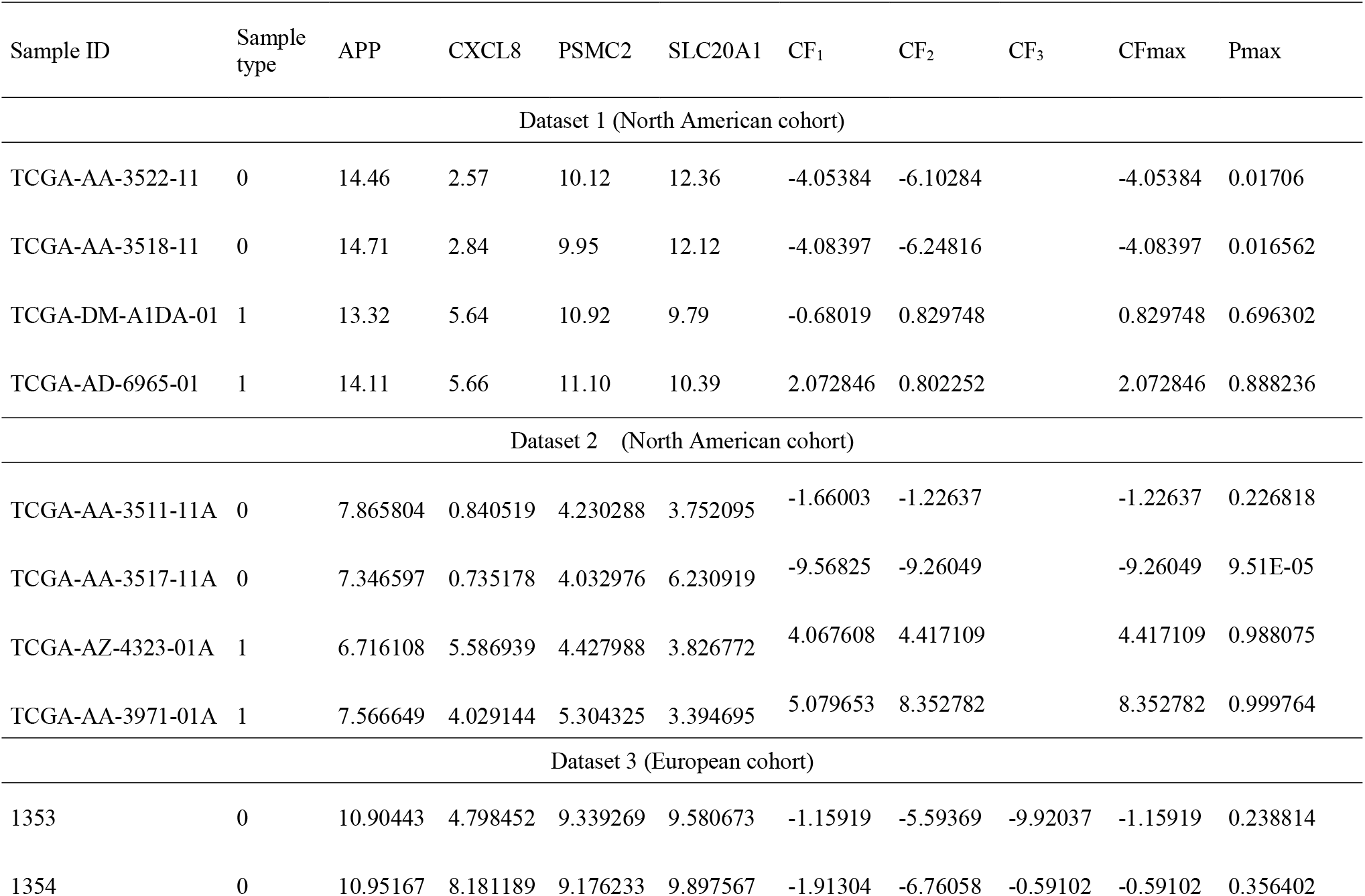
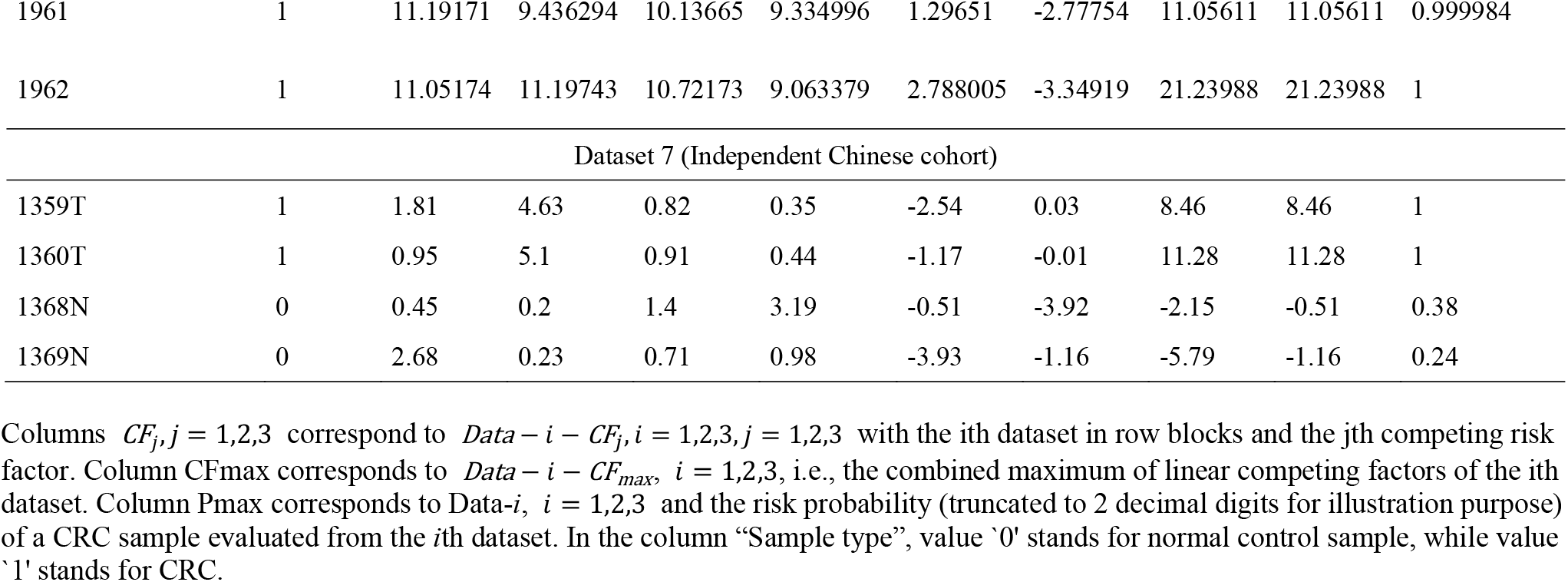
Gene expression values, competing factors, and risk probability in a portion of the samples

| Sample ID | Sample type | APP | CXCL8 | PSMC2 | SLC20A1 | CF<sub>1</sub> | CF<sub>2</sub> | CF<sub>3</sub> | CFmax | Pmax |
| --- | --- | --- | --- | --- | --- | --- | --- | --- | --- | --- |
| Dataset 1 (North American cohort) |  |  |  |  |  |  |  |  |  |  |
| TCGA-AA-3522-11 | 0 | 14.46 | 2.57 | 10.12 | 12.36 | -4.05384 | -6.10284 |  | -4.05384 | 0.01706 |
| TCGA-AA-3518-11 | 0 | 14.71 | 2.84 | 9.95 | 12.12 | -4.08397 | -6.24816 |  | -4.08397 | 0.016562 |
| TCGA-DM-A1DA-01 | 1 | 13.32 | 5.64 | 10.92 | 9.79 | -0.68019 | 0.829748 |  | 0.829748 | 0.696302 |
| TCGA-AD-6965-01 | 1 | 14.11 | 5.66 | 11.10 | 10.39 | 2.072846 | 0.802252 |  | 2.072846 | 0.888236 |
| Dataset 2 (North American cohort) |  |  |  |  |  |  |  |  |  |  |
| TCGA-AA-3511-11A | 0 | 7.865804 | 0.840519 | 4.230288 | 3.752095 | -1.66003 | -1.22637 |  | -1.22637 | 0.226818 |
| TCGA-AA-3517-11A | 0 | 7.346597 | 0.735178 | 4.032976 | 6.230919 | -9.56825 | -9.26049 |  | -9.26049 | 9.51E-05 |
| TCGA-AZ-4323-01A | 1 | 6.716108 | 5.586939 | 4.427988 | 3.826772 | 4.067608 | 4.417109 |  | 4.417109 | 0.988075 |
| TCGA-AA-3971-01A | 1 | 7.566649 | 4.029144 | 5.304325 | 3.394695 | 5.079653 | 8.352782 |  | 8.352782 | 0.999764 |
| Dataset 3 (European cohort) |  |  |  |  |  |  |  |  |  |  |
| 1353 | 0 | 10.90443 | 4.798452 | 9.339269 | 9.580673 | -1.15919 | -5.59369 | -9.92037 | -1.15919 | 0.238814 |
| 1354 | 0 | 10.95167 | 8.181189 | 9.176233 | 9.897567 | -1.91304 | -6.76058 | -0.59102 | -0.59102 | 0.356402 |
| 1961 | 1 | 11.19171 | 9.436294 | 10.13665 | 9.334996 | 1.29651 | -2.77754 | 11.05611 | 11.05611 | 0.999984 |
| 1962 | 1 | 11.05174 | 11.19743 | 10.72173 | 9.063379 | 2.788005 | -3.34919 | 21.23988 | 21.23988 | 1 |
| Dataset 7 (Independent Chinese cohort) |  |  |  |  |  |  |  |  |  |  |
| 1359T | 1 | 1.81 | 4.63 | 0.82 | 0.35 | -2.54 | 0.03 | 8.46 | 8.46 | 1 |
| 1360T | 1 | 0.95 | 5.1 | 0.91 | 0.44 | -1.17 | -0.01 | 11.28 | 11.28 | 1 |
| 1368N | 0 | 0.45 | 0.2 | 1.4 | 3.19 | -0.51 | -3.92 | -2.15 | -0.51 | 0.38 |
| 1369N | 0 | 2.68 | 0.23 | 0.71 | 0.98 | -3.93 | -1.16 | -5.79 | -1.16 | 0.24 |

Columns $CF_j, j = 1, 2, 3$ correspond to $Data - i - CF_j, i = 1, 2, 3, j = 1, 2, 3$ with the $i$ th dataset in row blocks and the $j$ th competing risk factor. Column CFmax corresponds to $Data - i - CF_{max}, i = 1, 2, 3$ , i.e., the combined maximum of linear competing factors of the $i$ th dataset. Column Pmax corresponds to $Data - i, i = 1, 2, 3$ and the risk probability (truncated to 2 decimal digits for illustration purpose) of a CRC sample evaluated from the $i$ th dataset. In the column “Sample type”, value ‘0’ stands for normal control sample, while value ‘1’ stands for CRC.

The risk probability of a sample with CRC can be calculated using the logistic function in Data_*i_CF_j_*. Table 4 shows subgroups defined by individual classifiers and their combinations. In the table, ‘only’ indicates the CRC patients identified by a specified classifier only. According to the classifiers, there were at least seven subgroups of CRC patients. For example, in the first dataset, 4 patients were detected by Data-3-CF1, 1 patient was detected by Data-3-CF2, 94 patients were detected by Data-3-CF3, and 60 patients were detected simultaneously by all three classifiers. The patients in each subgroup were different from other subgroups in each dataset as they may possess different genetic characteristics, implying different genetic determinations among groups.

**Table 4.** Subgroups of patients by individual classifiers and their combinations

| Dataset | CF<sub>1</sub> only | CF<sub>2</sub> only | CF<sub>3</sub> only | CF<sub>1&2</sub> only | CF<sub>1&3</sub> only | CF<sub>2&3</sub> only | All CF<sub>1</sub>,CF<sub>2</sub>,CF<sub>3</sub> |
| --- | --- | --- | --- | --- | --- | --- | --- |
| 1 | 7 | 39 | NA | 238 | NA | NA | NA |
| 2 | 5 | 10 | NA | 455 | NA | NA | NA |
| 3 | 4 | 1 | 94 | 1 | 399 | 6 | 60 |
| 7 | 2 | 4 | 23 | NA | NA | 12 | NA |

Since only one classifier was defined for datasets 4, 5, and 6, patients were not further subgrouped.

### Analysis of an independent Chinese cohort

We assessed the performance of the four critical DEGs (i.e., CXCL8, PSMC2, APP, and SLC20A1) identified in the above public datasets in a Chinese cohort that we collected at Sun Yat-sen University of Medical Sciences, China. Setting ***K*** = 1 and solving Equation (5), the classifiers were obtained (Table 2). It can be seen the classifier achieved an overall accuracy of 88.04%, sensitivity of 91.11%, and specificity of 85.11%. Interestingly, the coefficient of APP had a different direction (i.e., negative) in this Chinese cohort compared to other cohorts (North American, European, and Israel cohorts), implying diverse effects of this gene in various populations/ethnicities.

### Characterization of clinical and pathological features

To characterize the differences between subgroups defined by classifiers, we examined the attributes, including sex, age, histologic grades, and TNM tumor stages (Table 5). Since CRC patients in Datasets 4, 5, and 6 were identified by one single CF, the distribution of sample attributes was not shown. It should be noted that not all samples had complete clinical/pathological information. In the first and second datasets, the majority of CRC patients was determined by CF(1, 2). The numbers in the parentheses mean either CF1 and CF2 could determine the cancer status. There was no significant difference in gender preference, and the patients were identified in almost all age groups. In the second dataset, patients identified by individual CF1 or CF2 appeared to have late stage CRC, but the sample size in these subgroups were small. In the third dataset, CF3 appeared to be the classifier defined the majority of patients. There were no significant gender preference between groups. Stages 2 and 3 CRC were more common in subgroups associated with CF3.

**Table 5.** Distribution of clinical and pathological characteristics

The first dataset (North American cohort)
| Classifiers | Sex |  | Age at diagnosis (years) |  |  |  |  | TNM Stage |  |  |  |
| --- | --- | --- | --- | --- | --- | --- | --- | --- | --- | --- | --- |
|  | Male | Female | (20,50] | (50,60] | (60,70] | (70,80] | (80,100] | I | II | III | IV |
| CF<sub>1</sub> | 4 | 2 | 1 | 0 | 2 | 3 | 0 | 1 | 3 | 1 | 1 |
| CF<sub>2</sub> | 19 | 19 | 5 | 5 | 14 | 8 | 5 | 4 | 14 | 13 | 5 |
| CF<sub>(1,2)</sub> | 131 | 100 | 38 | 49 | 55 | 62 | 26 | 39 | 89 | 62 | 33 |

The second dataset (North American cohort)
| Classifiers | Sex |  | Age at diagnosis (years) |  |  |  |  | TNM Stage |  |  |  |
| --- | --- | --- | --- | --- | --- | --- | --- | --- | --- | --- | --- |
|  | Male | Female | (20,50] | (50,60] | (60,70] | (70,80] | (80,100] | I | II | III | IV |
| CF<sub>1</sub> | 1 | 1 | 1 | 0 | 1 | 0 | 0 | 0 | 0 | 2 | 0 |
| CF<sub>2</sub> | 4 | 4 | 2 | 2 | 3 | 0 | 1 | 0 | 0 | 8 | 0 |
| CF<sub>(1,2)</sub> | 229 | 198 | 55 | 73 | 109 | 126 | 62 | 73 | 167 | 115 | 61 |

The third dataset (European cohort)
| Classifiers | Sex |  | Age at diagnosis (years) |  |  |  |  | TNM Stage |  |  |  |
| --- | --- | --- | --- | --- | --- | --- | --- | --- | --- | --- | --- |
|  | Male | Female | (20,50] | (50,60] | (60,70] | (70,80] | (80,100] | I | II | III | IV |
| CF<sub>1</sub> | 3 | 1 | 1 | 2 | 0 | 1 | 0 | 0 | 1 | 2 | 1 |
| CF<sub>2</sub> | 0 | 1 | 0 | 1 | 0 | 0 | 0 | 0 | 1 | 0 | 0 |
| $CF_3$ | 47 | 47 | 11 | 13 | 27 | 31 | 12 | 9 | 44 | 33 | 8 |
| $CF_{(1,2)}$ | 1 | 0 | 0 | 0 | 1 | 0 | 0 | 0 | 1 | 0 | 0 |
| $CF_{(1,3)}$ | 222 | 177 | 55 | 58 | 114 | 122 | 49 | 20 | 184 | 153 | 41 |
| $CF_{(2,3)}$ | 4 | 2 | 0 | 1 | 3 | 1 | 1 | 0 | 4 | 2 | 0 |
| $CF_{(1,2,3)}$ | 33 | 27 | 9 | 6 | 20 | 15 | 10 | 4 | 28 | 15 | 10 |

The seventh dataset (Independent Chinese cohort)
| Classifiers | Sex |  | Age at diagnosis (years) |  |  |  |  | TNM Stage |  |  |  |
| --- | --- | --- | --- | --- | --- | --- | --- | --- | --- | --- | --- |
|  | Male | Female | (20,50] | (50,60] | (60,70] | (70,80] | (80,100] | I | II | III | IV |
| $CF_1$ | 0 | 2 | 0 | 1 | 0 | 0 | 1 | 0 | 2 | 0 | 0 |
| $CF_2$ | 1 | 3 | 1 | 1 | 1 | 0 | 1 | 2 | 0 | 0 | 1 |
| $CF_3$ | 13 | 10 | 5 | 3 | 3 | 8 | 4 | 0 | 7 | 11 | 5 |
| $CF_{(2,3)}$ | 8 | 4 | 0 | 1 | 6 | 3 | 2 | 1 | 7 | 2 | 2 |

### 3.2. Figures, Tables and Schemes

## 4. Discussion

It should be noted gene-gene interactions defined in our model are completely different from interaction effects widely used in experimental designs, such as row-column interaction effects or laboratory-chemical formula interaction effects in agriculture and industry. It is also different from the interaction term in linear regression analysis, i.e., using the multiplication of two covariates (predictors) to form an additional covariate to study the interaction effects of these two covariates in existing statistical models and machine learning. Using Dataset-1 as an example, CF1 and CF2 have different combinations of the four genes, with PSMC2 and SLC20A1 appearing twice. Their coefficient values were influenced by the other two genes in combination, clearly implying gene-gene and gene-subtype (CFi) interactions. Such interaction effects were also observed in Dataset-1-CF2, Dataset-2-CF2, and Dataset-3-CF3. Each classifier listed in Table 2 could a biomarker for CRC. Since both PSMC2 and CXCL8 were associated with CRC at different stages, the classifiers’ effects were likely modulated by these two major genes. Using a basketball team as an analog. These critical genes correspond to basketball players in a team. The team has three main teammates combinations for scoring. A positive coefficient associated with a player in a teammate scoring combination means that the longer the ball-controlling time by the player, the higher chance the team to score, and a negative coefficient associated with a player means that the shorter the ball-controlling time by the player, the higher chance the team to score. In the meantime, a question is which scoring combination is going to score. Under some scenarios, only one combination can score; under some scenarios, two of the three combinations can score; and under some scenarios, any combination can score. Figure 3 (Venn Diagram) clearly displays such phenomena. In the TCGA example, there are gene-gene interactions between competing factors, e.g., through PSMC2 and SLC20A1.

The functional relevance of the four critical DEGs to carcinogenesis has been reported. As a member of the CXC cytokine family, CXCL8 is one of the most significantly upregulated chemokines in CRC and contributes to tumor growth, invasion, and metastasis in patients [31–35, 36]. In addition, CXCL8 induces cell migration in colon cancer cells, acting as an autocrine growth factor [37, 38, 39]. Consistent with these observations, our analyses suggest CXCL8 could be a sensitive biomarker for early-stage CRC. PSMC2 is a key member of the 19S RP, a component of the 26S proteasome complex. PSMC2 has been implicated in the progression of ovarian cancer [40], pancreatic cancer [41], and osteosarcoma [42]. The 26S proteasome complex regulates many cellular processes, such as cell cycle progression, apoptosis, signal transduction, DNA repair and gene transcription, which are linked to cancer progression [43]. High expression of PSMC2 in CRC samples was associated with a poorer survival rate, and the anti-tumor effect of PSMC2 silencing was confirmed at the cellular level in vitro, suggesting silencing of PSMC2 could be a therapeutic strategy for CRC [44]. APP is involved in cell survival, cellular adhesion, differentiation, and migration [45]. The expression of APP was correlated with tumorigenesis and tumor growth of several human cancers [46, 47]. In our analyses, decreased APP expression was associated with reduced risk of CRC in the US, European, and Israel populations, which is consistent with the literature. However, a reverse effect was observed in Asian populations (Chinese and Japanese). The protein encoded by SLC20A1 is a sodium-phosphate symporter and plays a fundamental housekeeping role in supplying phosphate to cells [48, 49]. SLC20A1 gene knockout mice exhibit hypoplastic livers [50]. SLC20A1 is involved in tumor formation by HeLa cells in xenografted mice [51]. At present, the significance of SLC20A1 in CRC is largely unknown. In our analyses, coefficients of SLC20A1 have a negative sign in all the datasets, implying SLC20A1 expression is inhibited in CRC.

A strength of our method is that it works well for gene expression data from different platforms (RNA-seq and microarray) without a need to correct batch effects. Furthermore, the classifiers reached a high performance of cancer identification without correcting for tumor stages, ethnicities, and other clinical factors in the model as covariates. However, a limitation is that complete clinical and pathological information is not available from some public datasets. Particularly, analysis of prognostic value of the four DEGs is impossible given a lack of accurate clinical follow-up data. Nonetheless, the machine learning method adopted here is novel which has never been attempted before by other groups. Moreover, the four genes were validated in diverse populations/ethnicities. Therefore, this study is not a simple reanalysis of the public dataset but represents a pioneering effort in bringing in a novel concept (i.e., max-linear competing risk factor models) to the field of transcriptomic studies of human malignancies.

Our study has several limitations. First, this is a retrospective study analyzing large public genomic datasets. It would be ideal for performing prognosis analysis to assess the value of the critical genes in CRC. However, evaluation of the prognostic value of the classifiers is impossible due to the lack of complete clinical follow-up data. Nonetheless, this can certainly be done in future work as a nearly perfect performance of the four-gene-based classifiers provide an essential basis for designing prospective studies to assess the prognosis. Second, the current diagnosis of CRC is largely based on patients’ symptoms and clinical workups such as colonoscopies and tissue biopsies. Thus, it is difficult to find clinical significance in finding the critical DEGs identified in this study for CRC diagnosis. However, investigating molecular subtypes based on transcriptomic patterns is critical for discovering the underlying genetic mechanisms of carcinogenesis. Incorporating reliable genomic biomarkers such as the four-gene-based classifiers in the CRC diagnosis algorithm will further enhance the accuracy of disease identification and classification of patients and, eventually personalized medicine. Third, our study is based on gene expression profiles of CRC tissues. Whether these four-gene-based classifiers are applicable to the identification of CRC patients in the general population in blood samples awaits validation for future studies. We are planning on designing such a study in which the study cohorts have both CRC tissues and blood samples for analysis. Therefore, our study is not merely a reanalysis of the public data and identification of genes with known functions to CRC, but rather represents a pioneering effort in applying conceptually max-linear competing risk factor models to identify genomic/transcriptomic signatures for human malignancies.

A question is whether the identified critical DEGs could be chance findings. We acknowledge our results require further studies to validate. However, considering the accuracy in some cohorts is as high as 100%, the probability of chance finding is less than 10^-11^. We haven’t seen any other existing methods reported such reliable, robust, and interpretable findings across more than three cohorts, which reveals that our findings are unique and insightful. Moreover, we set very stringent criteria for defining critical DEGs in this work. Most of the four DEGs individually have known functional relevance to carcinogenesis, as proven in previous studies. Our work is the first to describe their interaction effects in determining the status of CRC. The findings could be a starting point for further work, such as gene network analysis, testing other related genes and their functional interaction, and discovering causal effects.

## 5. Conclusions

Genomic profiling studies play an important role in the identification of genes and biological pathways important for CRC. Currently, analysis of transcriptomic data suffers from several major limitations. First, the number of genes/transcripts analyzed is ultralarge compared to the number of study samples. Second, most studies have used conventional statistical methods, which suffer from limited power given a large number of genes/transcripts analyzed. Third, gene-gene interaction is not well addressed in the existing analytical models. Some of the DEGs could be noise effects or chance findings. Finally, it remains challenging to determine which genes are the key drivers of cancer development [28]. The new machine learning methods, as applied in this study, can partially overcome some of these limitations. Importantly, using the proposed smallest subset and the smallest number of signatures, DEGs can be identified in heterogeneous cohorts, even when covariates (i.e., expression values) are measured at different scales. Another unique feature is that the competing factors take into account gene-gene interactions within and between competing factors. In our early efforts, several critical DEGs were successfully identified for lung cancer [29], breast cancer [30], and COVID-19 [22,53–55] using the max-linear competing factor models. The present study further confirmed the power and validity of our method.

In this study, four critical DEGs were identified with high performance for the identification of CRC in seven datasets in different populations/ethnicities covering a spectrum of tumor stages and histologic grades. The four genes (i.e., CXCL8, PSMC2, APP, and SLC20A1) could describe the overall features of CRC at the genomic level. Such high performance has not been achieved in prior transcriptomic analyses of CRC. These four genes differ from those in our earlier work [29,30] and our ongoing research on liver and stomach cancers, and they can be concluded as CRC-specific and druggable and vaccinable targets.

## Supplementary Materials

Real data and computer outputs are in a supplementary file available online and submitted together with this paper. A MATLAB® demo code for solving an in Equation (4) (*λ*_2_ =0) is also available.

## Data Availability

The links of the public datasets are provided in Section “Data Description”. The dataset obtained from the independent Chinese cohort will be made available upon the request from readers.

## References

1. A. Jemal, F. Bray, M.M. Center, J. Ferlay, E. Ward, D. Forman, Global cancer statistics, CA Cancer J Clin, 61 (2011) 69–90.

2. N. Keum, E. Giovannucci, Global burden of colorectal cancer: emerging trends, risk factors and prevention strategies, Nat Rev Gastroenterol Hepatol, 16 (2019) 713–732.

3. B.K. Edwards, E. Ward, B.A. Kohler, C. Eheman, A.G. Zauber, R.N. Anderson, A. Jemal, M.J. Schymura, I. Lansdorp-Vogelaar, L.C. Seeff, M. van Ballegooijen, S.L. Goede, L.A. Ries, Annual report to the nation on the status of cancer, 1975-2006, featuring colorectal cancer trends and impact of interventions (risk factors, screening, and treatment) to reduce future rates, cancer, 116 (2010) 544–573.

4. P. Patel, P. De, Trends in colorectal cancer incidence and related lifestyle risk factors in 15-49-year-olds in Canada, 1969-2010, Cancer Epidemiol, 42 (2016) 90–100.

5. R.L. Siegel, S.A. Fedewa, W.F. Anderson, K.D. Miller, J. Ma, P.S. Rosenberg, A. Jemal, Colorectal Cancer Incidence Patterns in the United States, 1974–2013, J Natl Cancer Inst, 109 (2017).

6. J.P. Young, A.K. Win, C. Rosty, I. Flight, D. Roder, G.P. Young, O. Frank, G.K. Suthers, P.J. Hewett, A. Ruszkiewicz, E. Hauben, B.A. Adelstein, S. Parry, A. Townsend, J.E. Hardingham, T.J. Price, Rising incidence of early-onset colorectal cancer in Australia over two decades: report and review, J Gastroenterol Hepatol, 30 (2015) 6–13.

7. E.R. Fearon, Molecular genetics of colorectal cancer, Annu Rev Pathol, 6 (2011) 479–507.

8. R. Dienstmann, L. Vermeulen, J. Guinney, S. Kopetz, S. Tejpar, J. Tabernero, Consensus molecular subtypes and the evolution of precision medicine in colorectal cancer, Nat Rev Cancer, 17 (2017) 268.

9. A. Schlicker, G. Beran, C.M. Chresta, G. McWalter, A. Pritchard, S. Weston, S. Runswick, S. Davenport, K. Heathcote, D.A. Castro, G. Orphanides, T. French, L.F. Wessels, Subtypes of primary colorectal tumors correlate with response to targeted treatment in colorectal cell lines, BMC Med Genomics, 5 (2012) 66.

10. P. Roepman, A. Schlicker, J. Tabernero, I. Majewski, S. Tian, V. Moreno, M.H. Snel, C.M. Chresta, R. Rosenberg, U. Nitsche, T. Macarulla, G. Capella, R. Salazar, G. Orphanides, L.F. Wessels, R. Bernards, I.M. Simon, Colorectal cancer intrinsic subtypes predict chemotherapy benefit, deficient mismatch repair and epithelial-to-mesenchymal transition, Int J Cancer, 134 (2014) 552–562.

11. L. Marisa, A. de Reynies, A. Duval, J. Selves, M.P. Gaub, L. Vescovo, M.C. Etienne-Grimaldi, R. Schiappa, D. Guenot, M. Ayadi, S. Kirzin, M. Chazal, J.F. Flejou, D. Benchimol, A. Berger, A. Lagarde, E. Pencreach, F. Piard, D. Elias, Y. Parc, S. Olschwang, G. Milano, P. Laurent-Puig, V. Boige, Gene expression classification of colon cancer into molecular subtypes: characterization, validation, and prognostic value, PLoS Med, 10 (2013) e1001453.

12. A. Sadanandam, C.A. Lyssiotis, K. Homicsko, E.A. Collisson, W.J. Gibb, S. Wullschleger, L.C. Ostos, W.A. Lannon, C. Grotzinger, M. Del Rio, B. Lhermitte, A.B. Olshen, B. Wiedenmann, L.C. Cantley, J.W. Gray, D. Hanahan, A colorectal cancer classification system that associates cellular phenotype and responses to therapy, Nat Med, 19 (2013) 619–625.

13. E.M.F. De Sousa, X. Wang, M. Jansen, E. Fessler, A. Trinh, L.P. de Rooij, J.H. de Jong, O.J. de Boer, R. van Leersum, M.F. Bijlsma, H. Rodermond, M. van der Heijden, C.J. van Noesel, J.B. Tuynman, E. Dekker, F. Markowetz, J.P. Medema, L. Vermeulen, Poor-prognosis colon cancer is defined by a molecularly distinct subtype and develops from serrated precursor lesions, Nat Med, 19 (2013) 614–618.

14. H. Wang, X. Wang, L. Xu, J. Zhang, H. Cao, Analysis of the transcriptomic features of microsatellite instability subtype colon cancer, BMC Cancer, 19 (2019) 605.

15. E. Budinska, V. Popovici, S. Tejpar, G. D’Ario, N. Lapique, K.O. Sikora, A.F. Di Narzo, P. Yan, J.G. Hodgson, S. Weinrich, F. Bosman, A. Roth, M. Delorenzi, Gene expression patterns unveil a new level of molecular heterogeneity in colorectal cancer, J Pathol, 231 (2013) 63–76.

16. J. Guinney, R. Dienstmann, X. Wang, A. de Reynies, A. Schlicker, C. Soneson, L. Marisa, P. Roepman, G. Nyamundanda, P. Angelino, B.M. Bot, J.S. Morris, I.M. Simon, S. Gerster, E. Fessler, E.M.F. De Sousa, E. Missiaglia, H. Ramay, D. Barras, K. Homicsko, D. Maru, G.C. Manyam, B. Broom, V. Boige, B. Perez-Villamil, T. Laderas, R. Salazar, J.W. Gray, D. Hanahan, J. Tabernero, R. Bernards, S.H. Friend, P. Laurent-Puig, J.P. Medema, A. Sadanandam, L. Wessels, M. Delorenzi, S. Kopetz, L. Vermeulen, S. Tejpar, The consensus molecular subtypes of colorectal cancer, Nat Med, 21 (2015) 1350–1356.

17. U. Alon, N. Barkai, D.A. Notterman, K. Gish, S. Ybarra, D. Mack, A.J. Levine, Broad patterns of gene expression revealed by clustering analysis of tumor and normal colon tissues probed by oligonucleotide arrays, Proc Natl Acad Sci U S A, 96 (1999) 6745–6750.

18. E. Kim, S. Jung, W.S. Park, J.H. Lee, R. Shin, S.C. Heo, E.K. Choe, J.H. Lee, K. Kim, Y.J. Chai, Upregulation of SLC2A3 gene and prognosis in colorectal carcinoma: analysis of TCGA data, BMC Cancer, 19 (2019) 302.

19. R. Salazar, P. Roepman, G. Capella, V. Moreno, I. Simon, C. Dreezen, A. Lopez-Doriga, C. Santos, C. Marijnen, J. Westerga, S. Bruin, D. Kerr, P. Kuppen, C. van de Velde, H. Morreau, L. Van Velthuysen, A.M. Glas, L.J. Van’t Veer, R. Tollenaar, Gene expression signature to improve prognosis prediction of stage II and III colorectal cancer, J Clin Oncol, 29 (2011) 17–24.

20. Q. Cui, Y. Xu, Z. Zhang, V. Chan, Max-linear regression models with regularization, Journal of Econometrics, 222 (2021) 579–600.

21. Q. Cui,; Zhang, Z. Max-Linear Competing Factor Models. J. Bus. Econ. Stat. 2017, 36, 62–74. https://doi.org/10.1080/07350015.2015.1137761.

22. Z. Zhang. The Existence of At Least Three Genomic Signature Patterns and At Least Seven Subtypes of COVID-19 and the End of the Disease. Vaccines, 10, 761, 2022. https://doi.org/10.3390/vaccines10050761.

23. Y. Liu, N.S. Sethi, T. Hinoue, B.G. Schneider, A.D. Cherniack, F. Sanchez-Vega, J.A. Seoane, F. Farshidfar, R. Bowlby, M. Islam, J. Kim, W. Chatila, R. Akbani, R.S. Kanchi, C.S. Rabkin, J.E. Willis, K.K. Wang, S.J. McCall, L. Mishra, A.I. Ojesina, S. Bullman, C.S. Pedamallu, A.J. Lazar, R. Sakai, N. Cancer Genome Atlas Research, V. Thorsson, A.J. Bass, P.W. Laird, Comparative Molecular Analysis of Gastrointestinal Adenocarcinomas, Cancer Cell, 33 (2018) 721–735 e728.

24. Y. Hong, T. Downey, K.W. Eu, P.K. Koh, P.Y. Cheah, A ‘metastasis-prone’ signature for early-stage mismatch-repair proficient sporadic colorectal cancer patients and its implications for possible therapeutics, Clin Exp Metastasis, 27 (2010) 83–90.

25. T. Matsuyama, T. Ishikawa, K. Mogushi, T. Yoshida, S. Iida, H. Uetake, H. Mizushima, H. Tanaka, K. Sugihara, MUC12 mRNA expression is an independent marker of prognosis in stage II and stage III colorectal cancer, Int J Cancer, 127 (2010) 2292–2299.

26. M. Sheffer, M.D. Bacolod, O. Zuk, S.F. Giardina, H. Pincas, F. Barany, P.B. Paty, W.L. Gerald, D.A. Notterman, E. Domany, Association of survival and disease progression with chromosomal instability: a genomic exploration of colorectal cancer, Proc Natl Acad Sci U S A, 106 (2009) 7131–7136.

27. Alexander Malinowski, Martin Schlather, Zhengjun Zhang, Intrinsically weighted means and non-ergodic marked point processes, Ann Inst Stat Math, 68 (2016) 1–24.

28. J.P. de Magalhaes, Every gene can (and possibly will) be associated with cancer, Trends Genet, (2021).

29. Z. Zhang, Functional effects of four or fewer critical genes linked to lung cancers and new subtypes detected by a new machine learning classifier, Journal of Clinical Trials, 11 (2021).

30. Z. Zhang, Lift the veil of breast cancers using four or fewer critical genes. Cancer Informatics 2022, 21:1–11.

31. H. Terada, T. Urano, H. Konno, Association of interleukin-8 and plasminogen activator system in the progression of colorectal cancer, Eur Surg Res, 37 (2005) 166–172.

32. Y. Ning, H.J. Lenz, Targeting IL-8 in colorectal cancer, Expert Opin Ther Targets, 16 (2012) 491–497.

33. M. Baggiolini, B. Dewald, B. Moser, Human chemokines: an update, Annu Rev Immunol, 15 (1997) 675–705.

34. G. Galffy, K.A. Mohammed, P.A. Dowling, N. Nasreen, M.J. Ward, V.B. Antony, Interleukin 8: an autocrine growth factor for malignant mesothelioma, Cancer Res, 59 (1999) 367–371.

35. C. Rubie, V.O. Frick, S. Pfeil, M. Wagner, O. Kollmar, B. Kopp, S. Graber, B.M. Rau, M.K. Schilling, Correlation of IL-8 with induction, progression and metastatic potential of colorectal cancer, World J Gastroenterol, 13 (2007) 4996–5002.

36. Y. Itoh, T. Joh, S. Tanida, M. Sasaki, H. Kataoka, K. Itoh, T. Oshima, N. Ogasawara, S. Togawa, T. Wada, H. Kubota, Y. Mori, H. Ohara, T. Nomura, S. Higashiyama, M. Itoh, IL-8 promotes cell proliferation and migration through metalloproteinase-cleavage proHB-EGF in human colon carcinoma cells, Cytokine, 29 (2005) 275–282.

37. R. Brew, J.S. Erikson, D.C. West, A.R. Kinsella, J. Slavin, S.E. Christmas, Interleukin-8 as an autocrine growth factor for human colon carcinoma cells in vitro, Cytokine, 12 (2000) 78–85.

38. A.J. Wilson, K. Byron, P.R. Gibson, Interleukin-8 stimulates the migration of human colonic epithelial cells in vitro, Clin Sci (Lond), 97 (1999) 385–390.

39. W.J. Jin, J.M. Xu, W.L. Xu, D.H. Gu, P.W. Li, Diagnostic value of interleukin-8 in colorectal cancer: a case-control study and meta-analysis, World J Gastroenterol, 20 (2014) 16334–16342.

40. D. Nijhawan, T.I. Zack, Y. Ren, M.R. Strickland, R. Lamothe, S.E. Schumacher, A. Tsherniak, H.C. Besche, J. Rosenbluh, S. Shehata, G.S. Cowley, B.A. Weir, A.L. Goldberg, J.P. Mesirov, D.E. Root, S.N. Bhatia, R. Beroukhim, W.C. Hahn, Cancer vulnerabilities unveiled by genomic loss, Cell, 150 (2012) 842–854.

41. J. Qin, W. Wang, F. An, W. Huang, J. Ding, PSMC2 is Up-regulated in Pancreatic Cancer and Promotes Cancer Cell Proliferation and Inhibits Apoptosis, J Cancer, 10 (2019) 4939–4946.

42. M. Song, Y. Wang, Z. Zhang, S. Wang, PSMC2 is up-regulated in osteosarcoma and regulates osteosarcoma cell proliferation, apoptosis and migration, Oncotarget, 8 (2017) 933–953.

43. S. Frankland-Searby, S.R. Bhaumik, The 26S proteasome complex: an attractive target for cancer therapy, Biochim Biophys Acta, 1825 (2012) 64–76.

44. J. He, J. Xing, X. Yang, C. Zhang, Y. Zhang, H. Wang, X. Xu, H. Wang, Y. Cao, H. Xu, C. Zhang, C. Wang, E. Yu, Silencing of Proteasome 26S Subunit ATPase 2 Regulates Colorectal Cancer Cell Proliferation, Apoptosis, and Migration, Chemotherapy, 64 (2019) 146–154.

45. J.Y.S. Tsang, M.A. Lee, T.H. Chan, J. Li, Y.B. Ni, Y. Shao, S.K. Chan, S.Y. Cheungc, K.F. Lau, G.M.K. Tse, Proteolytic cleavage of amyloid precursor protein by ADAM10 mediates proliferation and migration in breast cancer, EBioMedicine, 38 (2018) 89–99.

46. K. Takagi, S. Ito, T. Miyazaki, Y. Miki, Y. Shibahara, T. Ishida, M. Watanabe, S. Inoue, H. Sasano, T. Suzuki, Amyloid precursor protein in human breast cancer: an androgen-induced gene associated with cell proliferation, Cancer Sci, 104 (2013) 1532–1538.

47. S. Lim, B.K. Yoo, H.S. Kim, H.L. Gilmore, Y. Lee, H.P. Lee, S.J. Kim, J. Letterio, H.G. Lee, Amyloid-beta precursor protein promotes cell proliferation and motility of advanced breast cancer, BMC Cancer, 14 (2014) 928.

48. S.V. Johann, J.J. Gibbons, B. O’Hara, GLVR1, a receptor for gibbon ape leukemia virus, is homologous to a phosphate permease of Neurospora crassa and is expressed at high levels in the brain and thymus, J Virol, 66 (1992) 1635–1640.

49. M.P. Kavanaugh, D.G. Miller, W. Zhang, W. Law, S.L. Kozak, D. Kabat, A.D. Miller, Cell-surface receptors for gibbon ape leukemia virus and amphotropic murine retrovirus are inducible sodium-dependent phosphate symporters, Proc Natl Acad Sci U S A, 91 (1994) 7071–7075.

50. L. Beck, C. Leroy, S. Beck-Cormier, A. Forand, C. Salaun, N. Paris, A. Bernier, P. Urena-Torres, D. Prie, M. Ollero, L. Coulombel, G. Friedlander, The phosphate transporter PiT1 (Slc20a1) revealed as a new essential gene for mouse liver development, PLoS One, 5 (2010) e9148.

51. L. Beck, C. Leroy, C. Salaun, G. Margall-Ducos, C. Desdouets, G. Friedlander, Identification of a novel function of PiT1 critical for cell proliferation and independent of its phosphate transport activity, J Biol Chem, 284 (2009) 31363–31374. Author 1, A.B. Title of Thesis. Level of Thesis, Degree-Granting University, Location of University, Date of Completion.

52. Brouwer-Visser J, Cheng WY, Bauer-Mehren A, Maisel D, Lechner K, Andersson E, Dudley JT, Milletti F. Regulatory T-cell Genes Drive Altered Immune Microenvironment in Adult Solid Cancers and Allow for Immune Contextual Patient Subtyping. Cancer Epidemiol Biomarkers Prev. 2018 Jan;27(1):103–112. doi: 10.1158/1055-9965.EPI-17-0461.

53. Zhang Z. Five critical genes related to seven COVID-19 subtypes: A data science discovery. Journal of Data Science, 19(1):142–150, 2021. https://doi.org/10.6339/21-JDS1005.

54. Zhang Z. Genomic Biomarker Heterogeneities Between SARS-CoV-2 and COVID-19. Vaccines 2022, 10(10), 1657; https://doi.org/10.3390/vaccines10101657

55. Zhang Z, Genomic Transcriptome Benefits and Potential Harms of COVID-19 Vaccines Indicated from Optimized Genomic Biomarkers. Vaccines 2022, 10(11), 1774; https://doi.org/10.3390/vaccines10111774

